# CytoBinning: immunological insights from multi-dimensional data

**DOI:** 10.1101/321893

**Authors:** Yang Shen, Benjamin Chaigne-Delalande, Richard W.J. Lee, Wolfgang Losert

## Abstract

New cytometric techniques continue to push the boundaries of multi-parameter quantitative data acquisition at the single-cell level particularly in immunology and medicine. Sophisticated analysis methods for such ever higher dimensional datasets are rapidly emerging, with advanced data representations and dimensional reduction approaches. However, these are not yet standardized and clinical scientists and cell biologists are not yet experienced in their interpretation. More fundamentally their range of statistical validity is not yet fully established. We therefore propose a new method for the automated and unbiased analysis of high-dimensional single cell datasets that is simple and robust, with the goal of reducing this complex information into a familiar 2D scatter plot representation that is of immediate utility to a range of biomedical and clinical settings. Using publicly available flow cytometry and mass cytometry datasets we demonstrate that this method (termed CytoBinning), recapitulates the results of traditional manual cytometric analyses and leads to new and testable hypotheses.

## Author Summary

The increasingly large number of measurements that can now be made simultaneously using cytometry platforms have created the impression that 2D scatter plots, which used to be the center stage of cytometry data analysis, don’t contain enough information. However, sophisticated methods that fully embrace large numbers of measurements are hampered by the difficulties of interpreting high-dimensional datasets and this limits their practical utility. CytoBinning fills the gap of complexity between conventional manual analysis and complex automated analysis to extract deep content in scatter plots which can be later cascaded into more complicated clustering or classification algorithms to obtain novel biological insights.

## Introduction

Cytometry is a multi-parameter single-cell measurement technique that is widely used in biological and clinical studies [1–6]. One of the main uses of flow cytometry, which has had a major impact across the fields of immunology and medicine, is to differentiate immune cells compositions among cell types or patients. Modern flow cytometers can routinely measure 15–20 cellular markers on millions of cells from dozens of samples in one experiment, and can sort cells into subpopulations based on those markers. Recently mass cytometry has expanded the number of markers that can be measured simultaneously to 100, though the technique is destructive to cells and does not allow for sorting. The conventional way of analyzing flow cytometry data uses a gating strategy which requires the manual selection of regions of interest (ROI) on sequential 2D scatterplots. This type of analysis is very labor intensive and inefficient for such large datasets and also suffers from subjectivity in both the sequence of 2D scatterplots and selection of thresholds (ROI) [3,4,7–10]. Therefore, as both the number of cells analyzed and the number of markers quantified for each cell have grown over the past decade, novel automated and unbiased analysis methods for flow cytometry data are emerging [11].

These novel analysis methods can be divided into two categories based on the problem they address: 1) methods trying to mimic and automatize the process of manual gating [12–18]; and 2) methods trying to identify cell populations using all markers simultaneously without prior biological knowledge [19–22]. Some cutting-edge approaches to automating manual gating, such as flowDensity [16], are very successful in re-identifying cell subsets that match with manually gated subsets in an automatic, reproducible way. However, gating (both manual and automatic) relies heavily on prior experience to inform the sequence of markers to gate. Furthermore, in gating, researchers must define the cell phenotypes to look for in advance of their analysis, hence hindering discovery of novel cell types and not tapping into the full potential of the acquired data. Gating methods also only explore a very limited portion of the total data space, though unsupervised methods have been published that enhance the efficiency of data usage, with the potential to reveal otherwise hidden differences between datasets [23]. Most unsupervised methods that allow novel cell type discovery aim to identify regions with high cell density in multi-dimensional space [19,21,23–30]. This assumes cells form distinct phenotypes and that only cells inside those relative high-density areas (peaks) are of importance. However, cells that are in between two high-density clusters (valleys) may also have potential biological significance [31]. Another limitation of clustering based methods is that concatenating different samples (which is a widely used strategy [28,32]) with potential batch effects can be problematic, hence limiting the meaningful combination datasets across institutions (which is very common in clinical trials). In addition, these clustering based methods require estimation of nearest neighbors in high-dimensional space which suffers from “curse of dimensionality” and may lead to misleading results [33]. As a result, people have been calling for the use of lower dimensional methods such as gating based on 2D scatterplots [34].

In this paper, we present a new method for analyzing cytometry data that utilizes such 2D scatter plots. Instead of gating, we dig deeper into the scatter plots mining the information that are largely bypassed by other methods. This method is useful for the majority of comparative studies that aim to elucidate the difference between two groups of samples. Our method, which we term CytoBinning, identifies the most information rich 2D scatter plots and extracts biological insights from them. We show that biologically relevant differences can be discovered from the pairs of markers identified with this approach. First, we introduce CytoBinning with a synthetic dataset, and then apply it to two public high-dimensional single cell datasets, a flow cytometry dataset comparing composition in immune cells between old and young healthy human donors [21], and a mass cytometry dataset analyzing the immune signature of eight types of human tissues [35].

## Results

We synthesized two point-patterns based on the expression of two virtual markers: maker A and marker B. Ten samples were generated for each point-pattern. The first point-pattern, called pattern A, consists of three point-clusters. Two large clusters each contain 5,000 points and a third relatively small cluster contains about 2,000 points. The three clusters are randomly sampled from Gaussian distributions that centered at point (0, 4), (0, –4) and (4, 0) with standard deviation 2, 2, and 1 respectively. The second point-pattern, called pattern B, also consists of three point-clusters. The two large clusters are generated in the same way as point-pattern A, however, the third smaller point-pattern only contains 200 to 500 points, sampled from a Gaussian distribution centered at point (−4, 6) with standard deviation 1 (Fig. S1).

## Percentile-based binning is a coarse-grained representation of point patterns

An example of percentile-based binning is shown in Fig. 1 using one synthetic sample with point-pattern A. Points inside the point-pattern were first binned into 3 levels based on the expression of marker A and B independently, each level containing one third of points. The 3 levels for marker A and B were then combined on a 2D scatter plot to form 9 sub-regions (these sub-regions are called boxes). The percentage of points in each box changes depending on the point-pattern. This binning method has been used as an alternative method to calculate mutual information (MI) in a robust and computationally efficient way [36]. MI is a measure of dependence between two random variables widely used in gene network inference [37] as a general measure of interdependency between genes. In our method, instead of summarizing the binning information into one number (MI), we used percentage of points in each box as a coarse-grained representation of point-patterns to obtain detailed information of point-patterns.

**Figure 1.**
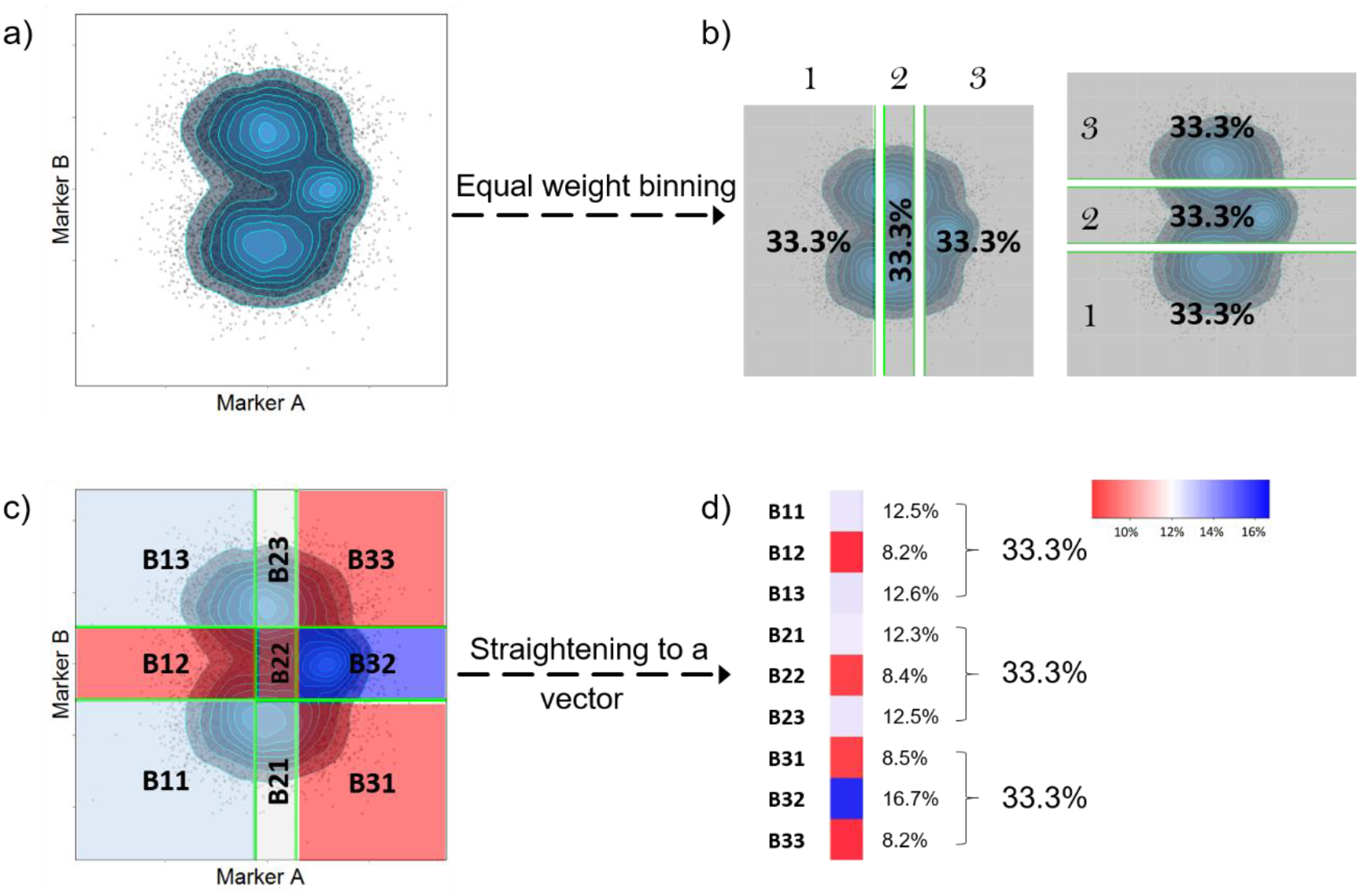
An example of percentile-based binning as a representation of 2D point-pattern (number of bins = 3). (**A**) Synthesized point-pattern formed by expression of marker A and marker B. (**B**) Points are binned into 3 bins each containing 1/3 (33.3 %) of the total points. The bins are labeled numerically based on the expression level of the related marker, with 1 the lowest and 3 the highest (similarly if points are divided into 5 bins then the highest level is 5). This binning is done independently for marker A and marker B based on their expression. (**C**) Bins obtained in (**B**) are combined so that 9 sub-regions referred as boxes are formed. The percentage of points inside each box is calculated, and the matrix of percentages is straightened to a vector so that the 2D point-pattern shown in panel (**A**) is coarse-grained to the vector in (**D**).

## Applying percentile-based binning to multiple samples enables meaningful classification

After we demonstrate how to represent point-patterns with percentile-based binning, next we show that this representation is able to capture real differences in point-patterns. Figure 2A shows two examples of the synthetic point-patterns. In total, 10 samples were generated for each point-pattern, and each sample was analyzed using percentile-based binning to generate the row vectors shown in Figure 2B. Using 3 bins, our method is able to cluster the two point-patterns into distinct groups, and correctly identifies the most significant difference (Fig. 2B & C). Boxplots of cell percentage in each box show that the box with most distinct difference between the two point-patterns is box B32 which contains the third cluster of point-pattern A (lower panel of Fig. 2B). This is the most significant difference between these two point-patterns, and it was captured without referring to density distribution of points. Minor differences between these two point-patterns (the small cluster located at the top left corner) were not spotted, since the percentage of points in box B13 is similar in both point patterns (Fig. 2B). However, this third cluster in pattern B (∼ 2 % to 5 %), was identified when the number of bins was increased to 6 (Fig. S2). Hence, the depth of analysis depends on the number of bins.

**Figure 2.**
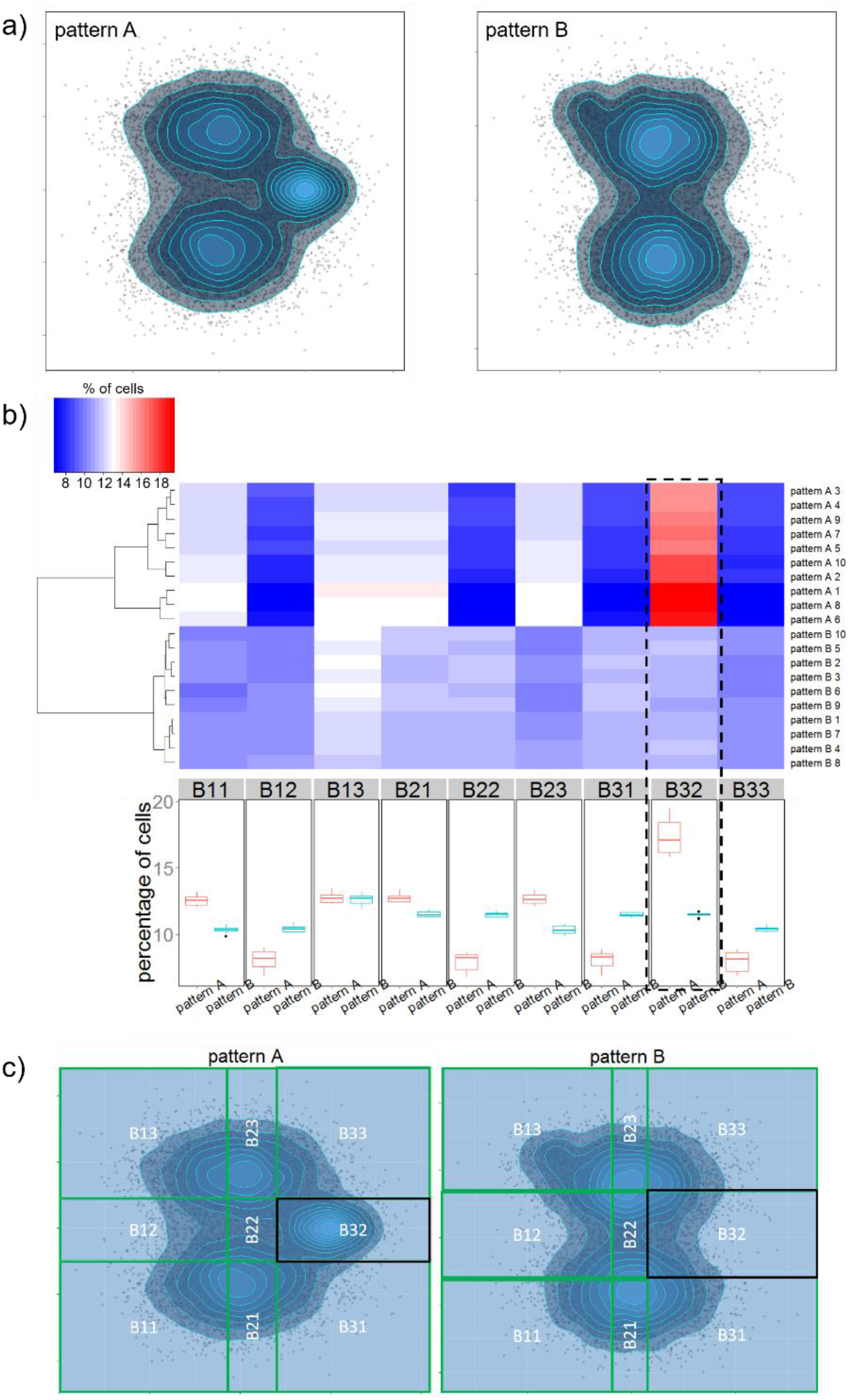
Percentile based binning is able to detect real difference between point-patterns. (**A**) Example of the two synthesized point-patterns A and B. The two large clusters in pattern A and B contain same number of cells and were generated with the same distribution. Pattern A contains a relatively large third cluster (10 % to 20 % of cells) at center right of the pattern and pattern B includes a smaller third cluster (2 % to 5 % of cells) on the top left corner. (**B**) Upper panel shows a heatmap of point percentage in each box for all samples, and lower panel shows boxplots of point percentage in each box between the groups of point-patterns. (**c**) Labels of each box. Highlighted is box B32.

The dotted black line represents FPR = 0.05. The number of bins that leads to a high FPR (> 0.05) is considered overfitting the dataset. (**B**) The maximum number of bins with FPR = 0 vs total number of samples used in the dataset. Blue dots show simulation results with our synthetic datasets and red dots shows the estimated number of bins using the rule of thumb: 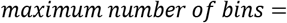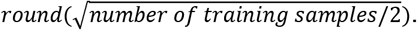.

## The maximum number of bins for binning depends on the number of samples (patients)

We’ve seen in the previous section that the depth of analysis depends on the number of bins used. And here we are going to show that the maximum number of bins we could use depends on the total number of samples (patients), for using a large number of bins to classify a small set of samples would cause overfitting. We see that false positive rate (FPR) increases with the numbers of bins used for binning (Fig. 3A). However, the maximum number of bins with tolerable FPR (FPR < 0.05) increased when we increase the number of samples from 20 to 60 (Fig. 3a). While with 20 samples we can only use as many as 3 bins to keep FPR under 0.05, with 60 samples this number increased to 6. And using 6 bins, our method is able to identify both of the differences we artificially generated between point-pattern A and B (Fig. S3). To get a general picture of how the maximum number of bins relates to number of samples, we calculated the maximum number of bins with FPR = 0 (we use this stringent condition because i) Synthetic data is easier to classify; ii) Real dataset contains more than 2 markers, and multiple tests correction should be taken into consideration) for various number of samples. We found that when the two groups to be classified contain the same number of samples (patients), the maximum number of bins is around the square root of half the sample size (Fig. 3B). In reality, the number of samples (patients) in different groups is rarely equal. However, we can overcome this inequality by assigning different number of samples to cross validation set for different groups so that in training dataset each group will have the same number of samples. Thus, once we know the number of samples in training dataset, we get a reasonable estimate for the number of bins to use.

**Figure 3.**
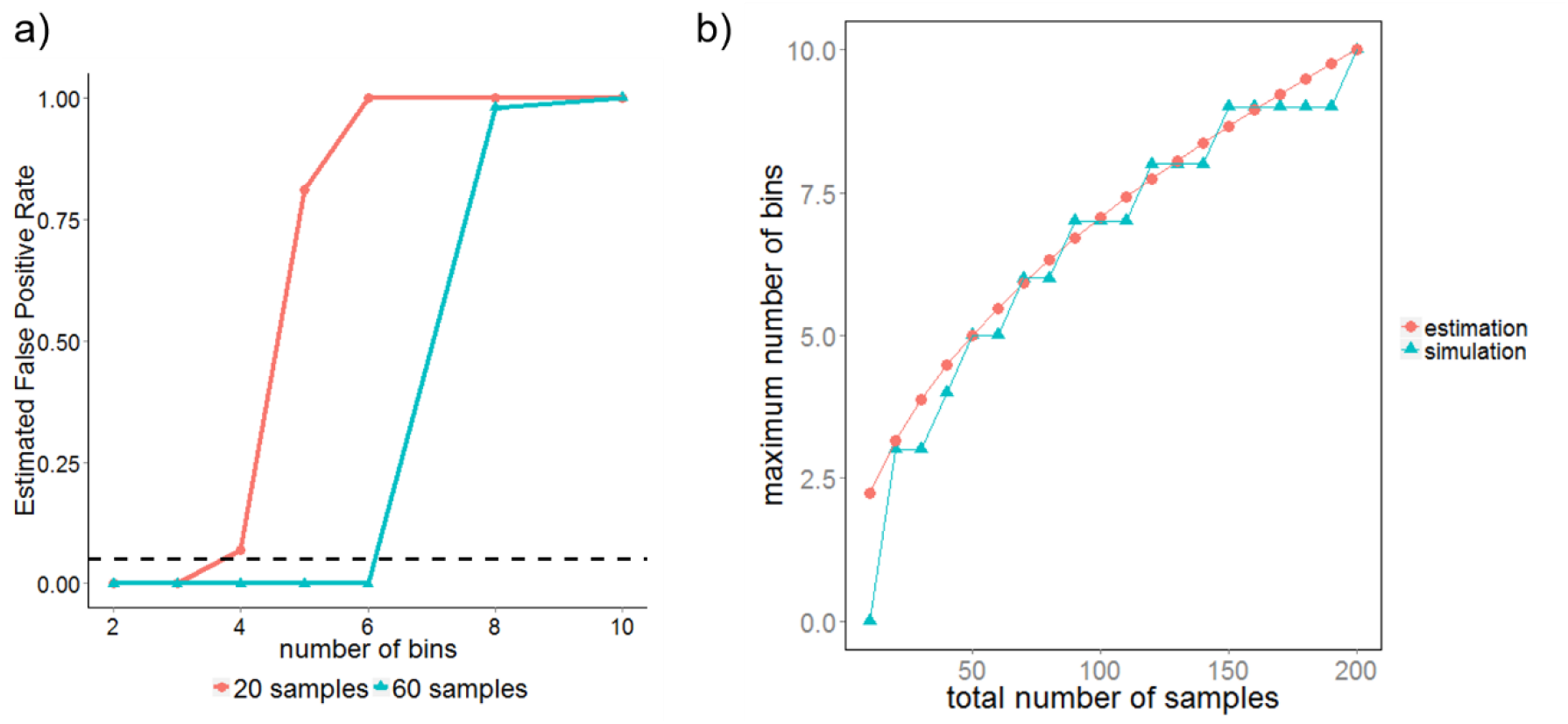
The maximum number of bins depends on the number of samples (patients). (**A**) Estimated false positive rate (FPR) vs number of bins for 20 samples (red) and 60 samples (blue).

## Application to two real human cytometry datasets

Next, we applied our method to two real flow cytometry datasets. Both datasets aim to identify differences between two biologically different patients/donor groups. In general, in order to get rid of debris and dead cells, some pre-processing steps should be taken before applying our method (e.g. manual/automatic gating to get live cells). In addition, depends on the question of interest, further gating can be applied to get more focused cell types, e.g. T cells, CD4^+^ T cells, etc. The pre-processed datasets are then the input for our method. We first determine the appropriate number of bins to use based on the number of samples in a dataset. Next, we apply the binning method showing in Figure 1 to the pre-processed dataset. Unlike the simulated dataset showing above which only contains two markers, real cytometry datasets usually measure much more markers which results in even more marker pairs. The binning method is applied to every possible pairs of markers. Then, in order to identify the important marker pairs, we separated the dataset into training and testing subsets. Using a classification algorithm called support vector machine (SVM) [38], we define important marker pairs as the ones that are able to achieve 100 % classification accuracy in both training and testing subsets. Once these marker pairs were determined, we move on to identify which regions formed by these marker pairs (boxes) are significantly different between the two groups.

### Old versus young

The first dataset we analyzed aims to find differences in the composition of immune cell types between old and young healthy donors [39]. Peripheral blood mononuclear cell (PBMC) samples from 34 healthy old donors (ages 60 and above) and 22 healthy young donors (ages 19 to 35) were taken, and their cellular composition were quantified by flow cytometry. In total, 16 markers were measured: Ki67, CD95, CD127, CD57, CD3, CD45RA, CD8, CD14, CCR4, CD27, CD11b, PD1, CD4, CD28, CCR7, and a viability dye (live/dead). We first manually gated for the live cells (Fig. S4) which were used as input for our method. At this stage, about 20 % of samples (4 young samples and 6 old samples) were randomly chosen as a cross validation set. We determined the optimal number of bins in remaining training dataset to be 5 (as the total number of samples in training set is 46, Fig. 3B), then we applied SVM classification based on the binning results of all possible pairs of markers. In total, we identified two pairs of markers (CD8 – CCR7, CD3 – CD4) that are able to classify old and young donors with 100 % accuracy both in training and testing dataset (Fig. S5). And boxes whose cell percentages are significantly different between old and young donors are identified. We selected the two boxes that are most different between old and young donors for demonstration below, remaining results can be found in supplementary information (Fig. S6 – S8).

### Naïve CD8^+^ T cells are found significantly decreased in elderly donors using only CD8 and CCR7 expression

We first look at box B55 which contains cells whose expression of both CD8 and CCR7 are in the top 20 % (i.e. CD8^high^ CCR7^high^, Fig. 4A). We find that percentage of cells inside box B55 decrease significantly in old donors (Fig. 4B). On the other hand, mean fluorescence intensity (MFI) of cells inside box B55 are similar among donors for all markers, indicating cells inside box B55 are homogeneous across all samples (Fig. 4C). Notice that CD3 and CD45RA MFI levels are high for all samples, and since cells inside box B55 already express highest 20 % of both CD8 and CCR7, one possibility is that cells inside B55 are naïve CD8^+^ T cells. Indeed, cells in B55 agrees well with manually gated naïve CD8 cells (Fig. S9 & S10A) on single cell level. In addition, when comparing the expression of CD45RA and CCR7 between cells in B55 and manually gated CD8 naïve and memory cell types we find that cells in B55 match well with naïve cells for young donors with slightly higher variation on CD45RA (Fig. 4D). Cells in B55 express higher variation in CD45RA for older donors, which is expected since box B55 was selected without expression information of CD45RA (Fig. 5E). Together, these results suggest that cells inside box B55 resemble naïve CD8^+^ T cells. Decreasing of naïve CD8^+^ T cells with ageing is a well-known observation in immunology [40] and is also identified in this dataset (Fig. S10B). In addition, we found that the abundancy of effector memory (TEM) and effector memory RA^+^ (TEMRA) CD8^+^ T cells are increased in old donors, as suggested by the increased percentage of cells in B51 (CD8^high^ CCR7^low^) (Fig. S11).

**Figure 4.**
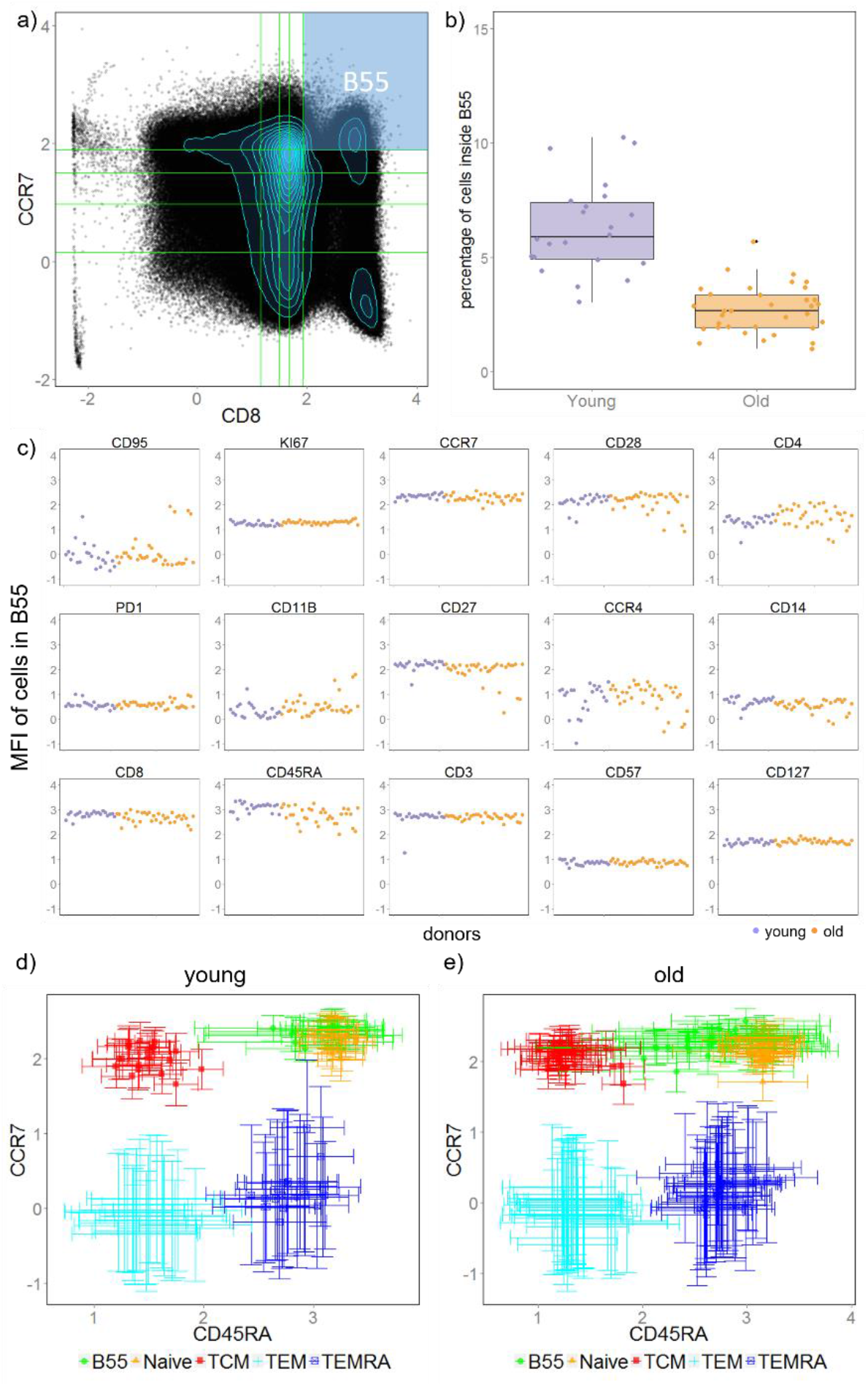
Naïve CD8^+^ T cells were identified by our method as significantly decreased in old donors using only two markers: CD8 and CCR7. (**A**) An example showing scatter plot of CD8 vs. CCR7 with box B55 highlighted. (**B**) Boxplot of cell percentage inside box B55 between the two groups of donors. Each dot is a donor. (**C**) Scatter plot of mean fluorescent intensity (MFI) of each donor, each point shows a donor (purple: young, orange: old). (**D**) & (**E**) MFI of CD45RA vs. MFI of CCR7 for cells in B51, naïve and memory CD8 T cells. Each symbol shows a donor (young donors in **D** and old donors in **E**), vertical and horizontal error bars show standard deviation of CCR7 and CD45RA intensity respectively.

### Distinction between naïve and memory CD8^+^ T cells is blurred in old donors

Next, we analyzed cells inside box B52 (CD8^high^CCR7^intermediate/low^). The percentage of cells inside box B52 (Fig. 5A) was found to be increased in the old group (Fig. 5B). Similar to box B55, the MFI of cells in box B52 for all samples were at similar levels for most markers, indicating a homogeneous cell subset is identified among all donors (Fig. 5C). Notice that box B52 lies in between two peaks (Fig. 5A) which is a region often neglected or assigned to one of the peaks by manual gating, and we have shown above that cells in peak above B52 (i.e. B55) resemble naïve CD8 T cells and cells in peak below B52 (i.e. B51) resemble memory CD8 T cells (TEM and TEMRA). We hence infer that cells in B52 are transition cells between naïve and memory cells which increases with ageing. Fig. 5D & E show how cells in B52 locate relative to manually gated naïve and memory CD8 T cells.

**Figure 5.**
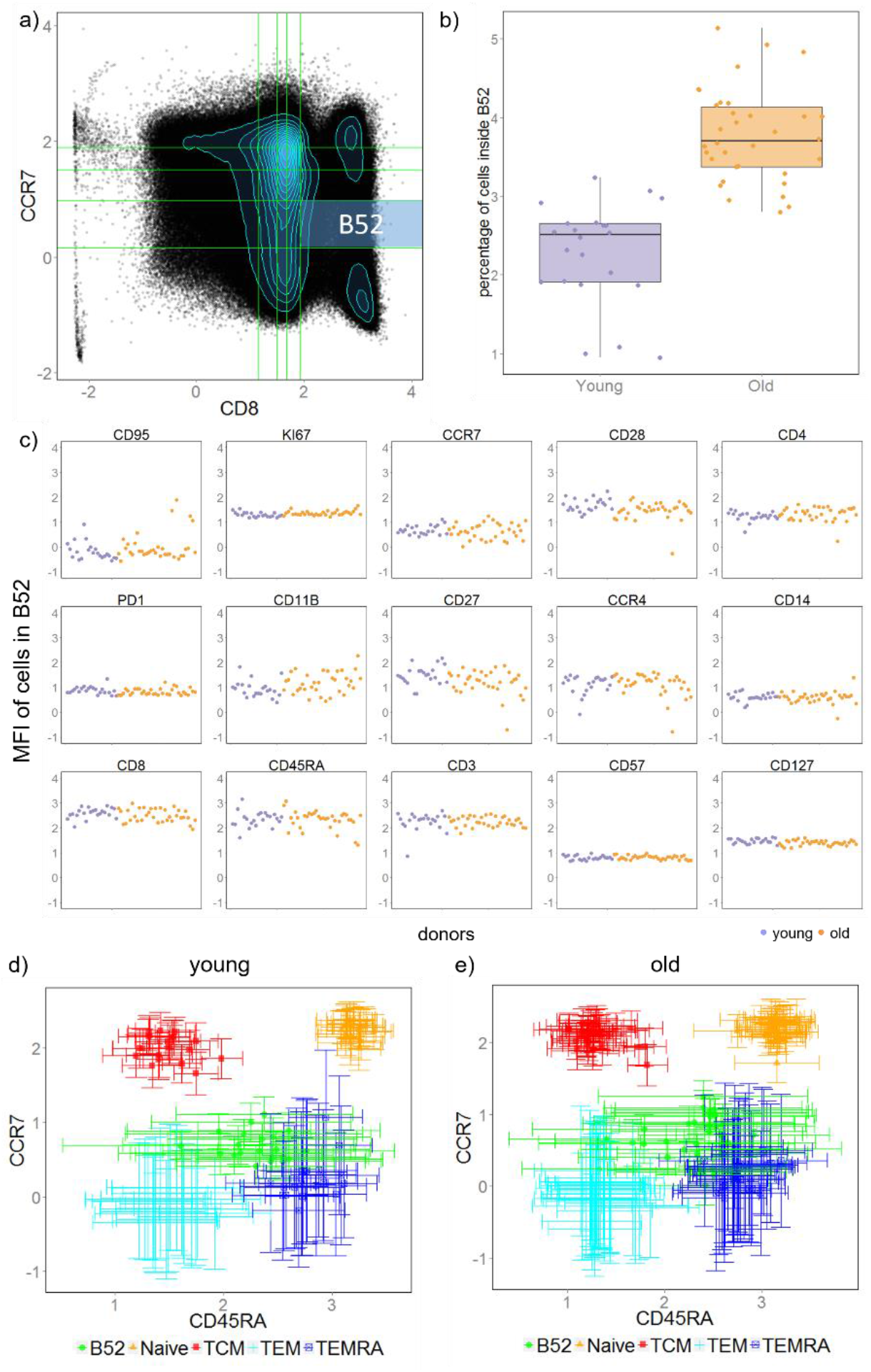
An intermediate cell region which is often neglected by gating methods is identified as significantly increased in old donors. (**A**) An example of scatter plot of CD8 vs. CCR7 with box B52 highlighted. (**B**) Boxplot of cell percentage inside box B52 between the two groups of donors. Each dot is a donor. (**C**) Scatter plot of mean fluorescent intensity (MFI) of each donor, each point shows a donor (purple: young, orange: old). (**D**) & (**E**) MFI of CD45RA vs MFI of CCR7 for cells in B52, naïve, and memory CD8 T cells. Each symbol shows a donor (young donors in **D** and old donors in **E**), vertical and horizontal error bars show standard deviation of CCR7 and CD45RA intensity respectively.

### CD4 versus CD8

Next, we applied our method to a mass cytometry dataset that originally aims to identify immune signatures among 8 types of human tissues: cord blood, PBMC, liver, spleen, skin, lung, tonsil and colon [35]. There are in total 35 samples, 3 to 6 samples for each type of tissue (see Methods). The marker panel used for mass cytometry contains 41 markers with a focus on the function (cytokine expression) of T cells (a full list of all 41 markers can be found in supplementary information and [35]). Instead of differentiating the 8 types of tissues, here we tried to classify CD4^+^ cells from CD8^+^ cells in all types of tissues. This is a good test for our method since there exists great within-group variance (different tissues) in the two groups we’re comparing, and we aim to find patterns that are consistent / similar across all types of tissues but are significantly different between CD4^+^ and CD8^+^ cells. Like the previous dataset, we divide these tissue samples into training and testing sets as well. From the 35 CD4 samples, 5 samples are randomly selected to be cross validation set; and the same was done for the 35 CD8 samples separately. Since there are in total 60 samples in training set (30 for each cell type), the number of bins to use is 5 (Fig. 3B). We identified 7 pairs of markers that were able to classify CD4^+^ and CD8^+^ cells with 100 % accuracy for both training and cross validation datasets. Only 1 marker pair (CCR10 vs. CCR9) out of the 7 contains purely trafficking markers. This indicates that CD4^+^ and CD8^+^ T cells can be more easily differentiated by their function and lineage markers than trafficking markers, which is consistent with the results in the original paper [35]. We selected one of the seven marker pairs: Interleukin (IL)-2 vs. CD25 to show in Figure 6. The pattern formed by CD4 cells is distinct from CD8 cells in that CD4 cells express significantly more IL-2 and slightly more CD25 in all types of tissues, which agrees with previous findings based on circulating immune cells [41]. In addition, we found that percentage of cells in box B13 (red shaded region in Figure 6, IL-2 low and CD25 intermediate) is significantly higher in CD8 cells, which is a subtle difference that would be missed by algorithms based on a peak finding.

**Figure 6.**
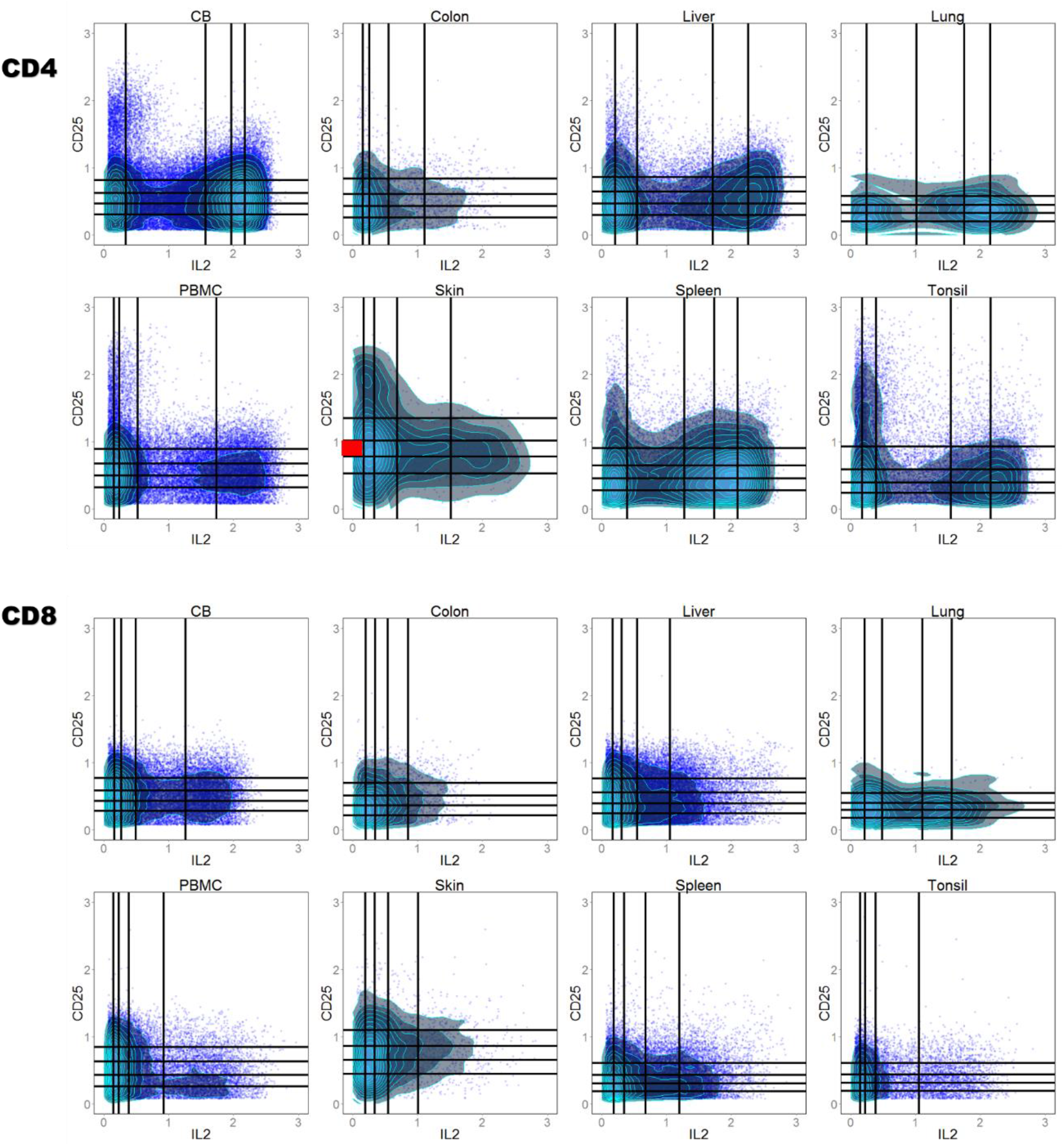
Patterns formed by IL-2 vs CD25 is distinct between CD4 and CD8 cells. One randomly chosen sample for each tissue is shown. The same sample for each type of tissue is chosen to illustrate both CD4 and CD8 cells. Percentage of cells in the red shaded box (B13: IL-2 negative and CD25 intermediate) is significantly higher in CD8 cells comparing to CD4 cells. Cells inside box B13 also express CD45RA, TNF, and CD127 (Fig. S12).

## Discussion

The complexity of cytometry data has increased significantly in the last few years due to the advancement in experimental techniques that enable measurements of dozens of parameters on each cell for millions of cells [9]. Novel analysis algorithms are being introduced at a rapid pace to deal with this data deluge that identify clusters of cells and project the high dimensional information graphically in innovative ways. However, these graphics are not directly interpretable and translatable into hypotheses and actions by biomedical researchers and clinicians. There is also the flaw that nearest neighbors are not meaningful in high dimensions, which is a phenomenon referred to as the “curse of dimensionality” [33,34]. Here we introduce a simpler, alternative approach we term CytoBinning. Our analysis approach combines automation of a more traditional workflow (as advocated in [34]) and machine learning which links the high dimensional data back to two biomarkers which can be represented as 2D scatter plots. The 2D scatter plot outputs are designed to be directly interpretable by biomedical researchers and clinicians, who have an established intuition for the meaning of these graphics. Thus, we are able to leverage their existing expertise in interpreting these kinds of scatterplots. When the differences in phenotype are small, CytoBinning is able to further focus the researcher or clinician’s attention by identifying, which specific regions of the scatter plot exhibits the most notable differences between two groups of donors, allowing subtle shifts in the immune phenotype to be highlighted.

In contrast to automated gating methods that focus on the exact position of density peaks or the number of groups formed by cells, CytoBinning doesn’t estimate the probability density distribution of cells, and thus its findings are not limited to regions with high cell density or sensitive to shifts in calibration. Instead, it extracts the pattern of 2D dot-plots and represents it with a sequence of cell percentages. This enables the comparison across samples measured in different experiments (given the markers are the same and they are measured in the same channel respectively). In addition, CytoBinning doesn’t require any *a priori* biological understanding to guide the path of analysis. Conversely, it provides a list of important marker pairs and related important cell sub-regions for biological researchers to subsequently interrogate.

In the first public dataset we analyzed, which compares lymphocyte populations in old and young healthy donors, CytoBinning automatically discovered a decrease of naïve CD8^+^ T cells in the elderly, a well-known yet subtle phenotype. In addition, CytoBinning identified a region in the scatterplot of relatively low cell density between two well-established cell clusters which is clearly increased with ageing as a new area of interest for the biological researcher. Two markers (CD8 and CCR7) are sufficient to pinpoint this subset of cells which resides between naïve and memory CD8^+^ T cells, and is not associated with a local peak in cell density in the scatterplot. Such an area would be missed by both manual gating and density-based algorithms, or by focusing exclusively on peaks in density.

The second public dataset we analyzed was even higher dimensional, based on mass cytometry from eight types of human tissues. CytoBinning analysis of CD4^+^ vs. CD8^+^ T cells automatically discovered higher expression of IL-2 in CD4^+^ T cells as we would expect [41], and shows that this overexpression is consistent throughout all eight types of human tissues studied. In addition, CytoBinning correctly identified that CD25 is also more highly expressed in CD4^+^ T cells [41]. This difference in CD25 and IL-2 was consistent among all types of tissues, which is known and therefore obvious to a biological researcher. However, it also demonstrates the power of our method as this marker pair was re-discovered without prior knowledge from a heterogeneous dataset incorporating 35 samples from 8 different tissues, each labelled with 41 markers. Hence, in addition to avoiding the pitfalls of density-based approaches, when applied to very high-dimensional datasets CytoBinning is able to select the salient markers which discriminate between groups of samples.

In summary, CytoBinning as a robust, automated approach to analyze high throughput cytometry data presented in familiar and interpretable 2D scatter plots. While simultaneous assessment of all markers is an important vision and challenge, in the interim there is a need to facilitate interpretation of high-dimensional data given the evident gap between our technological ability to acquire this information and our ability to understand it. CytoBinning fills the void between conventional manual analysis and complex automated analysis to extract deep content in scatterplots which can be later cascaded into more complicated clustering or classification algorithms to obtain novel biological insights. This has particular potential value in clinical and biological research settings where high-dimensional data is increasingly available and commonly not fully understood. CytoBinning is able to identify the most important markers, while also highlighting novel cell populations that distinguish comparator datasets even if these are to be found in areas of low cell density. Hence, it is a practical analysis approach with potential to fill the complexity gap in interpretation of high-dimensional data in a wide range of biomedical and clinical settings.

## Methods

### Binning

The binning we used in our method has been previously proposed to estimate mutual information (MI) [36]. Given bin number *b*, equally populated bins are drawn based on single cell expression of marker A and marker B independently. These bins are then overlaid on each other so that a grid is formed with *b* ^2^ regions (boxes). Percentage of cells inside each box is then an estimation of the joint probability *P(A_i_, B_j_)*, where *i* and *j* are the corresponding bins this box locates at. For a random distribution where marker A and marker B is not correlated in any way, *P(A_i_, B_j_)* should be approximately the same in for every box. This is not true if marker A and marker B is related in any way (i.e. their mutual information is not zero, this relationship can be both linear and nonlinear). We use all *P(A_i_, B_j_)*s as a coarse-grained representation of the point pattern between single cell expression of marker A and marker B. (Fig. 1) In our method this binning is done for every pair of markers.

### Determine appropriate number of bins

We deduced a relationship between the maximum number of bins with zero false positive rate (FPR) and the number of samples used in classification using our synthetic data. The relationship we found is: 
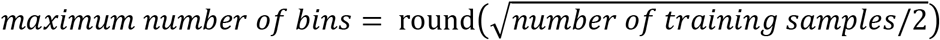

Thus, for a given dataset, an estimation of the number of bins to be used is achieved. In addition, we estimate FPR as follows:

1. For a given number of bins, apply afore-mentioned binning method to one pair of markers. Each sample is now represented by the vector of *P(A_i_, B_j_)*.
2. Randomly divide all samples into two groups.
3. Apply SVM classification (ksvm function in R package ks, with linear kernel and C = 10) on the randomly divided groups.
4. Repeat step 2 & 3 for 100 iterations, record the frequency when classification accuracy achieved 100 % in step 3.
5. Repeat step 1 to 4 for all marker pairs, calculate the mean frequency of one pair achieving 100 % accuracy. This frequency is used as an estimation of FPR.
6. Repeat steps above for all numbers of bins.

### Log ratio transformation

The percentages of cells in each box obtained with CytoBinning is compositional as they add up to 100. To get rid of this dependency, we divide the percentages by their median before taking log with base 2 for every sample and every marker pair.

### Selecting important marker pairs

Once the number of bins is determined, we divide all samples into training set (about 80 % of total samples) and testing set (the remaining 20 % of all samples). SVM is applied to training set and classification boundary obtained for every pair of markers. We use the obtained classification boundary to predict the cross validation set. Pairs that reached 100 % accuracy for both training and cross validation datasets are chosen as important marker pairs.

### Selecting important boxes

We combined boxes formed by all selected marker pairs and applied statistical test (wilcox) for percentage of cells in each box. We then corrected the p values for multiple comparison with Bonferroni correction, and boxes with p value <0.001 after correction are selected as important boxes. Important marker pairs selected above without any important boxes are eliminated from the important marker pair list.

### Dataset 1: Comparing old and young healthy PBMCs

**Overview of samples**. This dataset is published in reference [21] and downloaded at Flow Repository (http://flowrepository.org) website [42]. These samples were processed in two experiments, with 19 samples from young donors and 20 samples from old donors processed in the first experiment, and the remaining samples processed in the second experiment. The panel of markers were kept the same for both experiments. In total, 16 markers are measured: Ki67, CD95, CD127, CD57, CD3, CD45RA, CD8, CD14, CCR4, CD27, CD11b, PD-1, CD4, CD28, CCR7 and a viability dye (live/dead). Details of sample storage and processing can be found in [21].

**Pre-processing**. Downloaded FACS files were first compensated based on the spill matrix in the fcs files, and then manually gated to get live cells (Fig. S4). Logicle transformation was performed with w = 0.5, t = 262144, and m = 4.5 using logicleTransform function in flowCore package with R.

### Dataset 2: Comparing CD4 and CD8 T cells in various types of tissues

The dataset used for demonstration was first published in [35] and downloaded from flow repository website (https://flowrepository.org/) [42]. Tissue types, number of samples, and the reason for surgery are listed in Table 1. Immune cells were isolated from collected tissues and cryopreserved. They were then thawed and washed for mass cytometry experiment. Two panels of antibodies were used for staining, each containing 41 markers. The two panels were named as “Function” and “Traffic” according to the antibodies included in it. We only used function panel in this paper. Details of experimental process and the lists of antibodies can be found in [35]. The downloaded samples from flow repository are FACS files, pre-gated to major immune types (e.g. CD4, CD8, NKT, etc.). We used only CD4 and CD8 cells. We performed logicle transformation using logicleTransform in R package flowCore, with parameters w = 0.25, t = 16409, m = 4.5, and a = 0 according to [35]. The logicle transformed data were then saved as text files for further analysis.

**Table 1.**
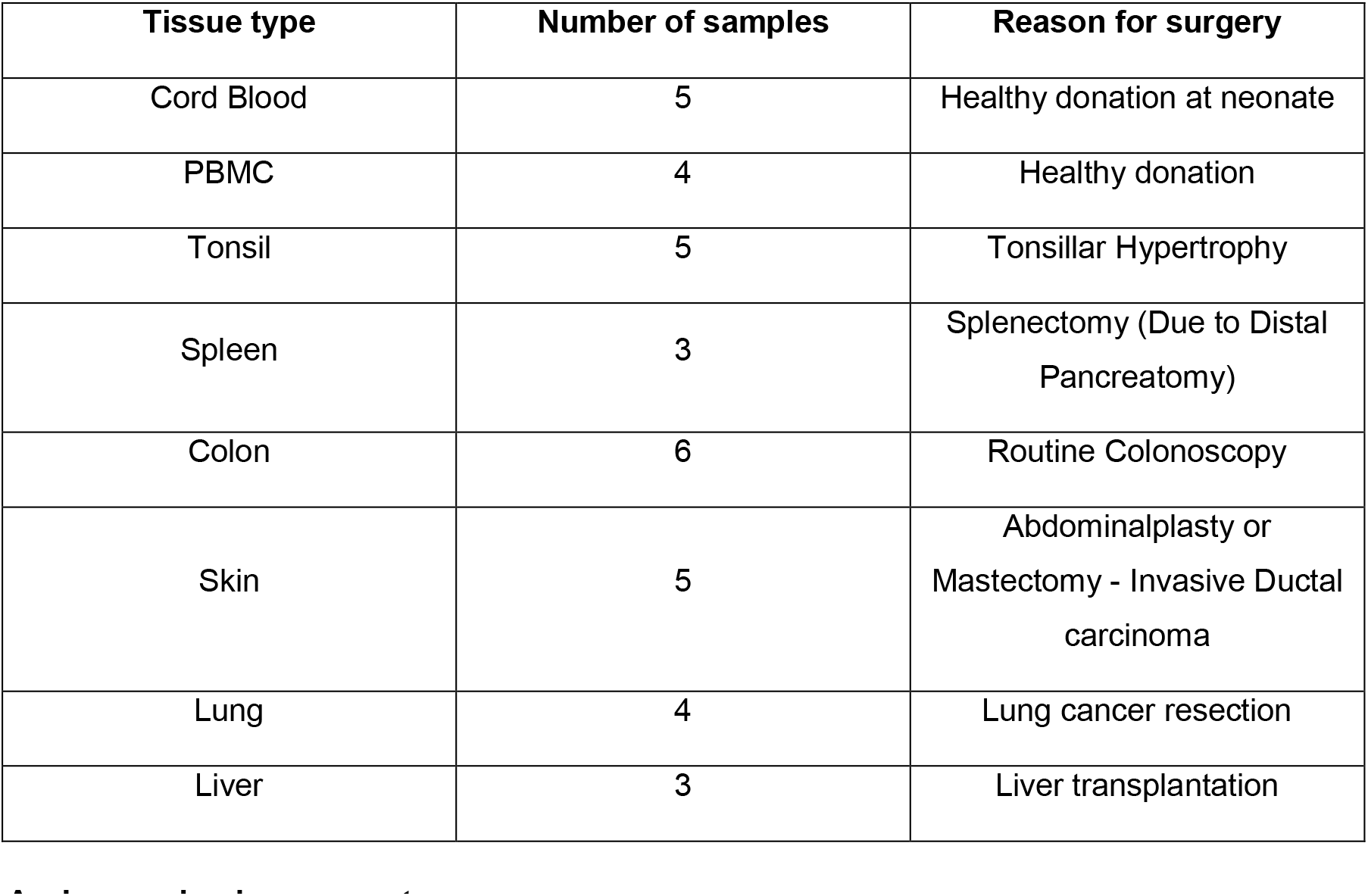
Summary of sample information

| Tissue type | Number of samples | Reason for surgery |
| --- | --- | --- |
| Cord Blood | 5 | Healthy donation at neonate |
| PBMC | 4 | Healthy donation |
| Tonsil | 5 | Tonsillar Hypertrophy |
| Spleen | 3 | Splenectomy (Due to Distal Pancreatomy) |
| Colon | 6 | Routine Colonoscopy |
| Skin | 5 | Abdominalplasty or Mastectomy - Invasive Ductal carcinoma |
| Lung | 4 | Lung cancer resection |
| Liver | 3 | Liver transplantation |

## Acknowledgements

R.W.J.L. was supported by the National Institute for Health Research (NIHR) Biomedical Research Centre based at Moorfields Eye Hospital NHS Foundation Trust and UCL Institute of Ophthalmology, UK. The views expressed are those of the authors and not necessarily those of the National Health Service, the NIHR or the Department of Health. W.L. was partially supported by AFOSR grant FA9550–16-1–0052. Y.S. and B.C.D. were supported by the National Institutes of Health, National Eye Institute intramural research program. The content of this publication does not necessarily reflect the views or policies of the U.S. Department of Health and Human Services, nor does mention of trade names, commercial products, or organizations imply endorsement by the U.S. Government.

## Supporting Information

**S1 Fig. Scatter plots of simulated point patterns.** First two rows show point pattern A, the lower two rows show point pattern B. Two major clusters in both point pattern A and B are generated from the same distributions. The third cluster of point pattern A, located on center right, consists about 10 to 20 % of total cells. The third cluster of point pattern B, located at upper left of all points, contains only 2 to 5 % of all cells.

**S2 Fig. Heatmap for percentage of cells inside each boxes with 6 bins.** Percentage of cells in box B16 (which corresponds to the third cluster in point pattern B) is significantly different between these two point patterns. This is not seen with only 3 bins. However, with 20 samples, analysis results using 6 bins is not reliable. Hence, in order to identify fine difference, more samples are needed.

**S3 Fig. With 6 bins, both differences between pattern A and pattern B can be found by CytoBinning.** a) Example for both pattern A and pattern B. b) Heatmap showing hierarchical clustering for CytoBinning results with 6 bins. Highlighted are the most different boxes between pattern A and B.

**S4 Fig. Illustration of manual gating strategy to get live cells.**

**S5 Fig. Select important marker pairs for the first dataset (old vs young).** Ten samples are randomly selected as cross validation dataset (4 in young group and 6 in old group). SVM classification was used to separate old and young samples with binning results for each marker pair separately. Two marker pairs are able to achieve 100 % classification accuracy for both trainning and cross validation dataset (CD4 vs CD3 and CD8 vs CCR7).

**Fig. 1. Ilustration of box B25 formed by CD4 and CD3.** a) Position of box B25. b) Percentage of cells in B25 is higher in young donors. c) Scatter plot of mean flourescent intensity (MFI) for all donors and all markers. This suggests cells in B25 are CD3+, CD8+ and CD45RA+. d) An example showing how cells in B25 (green) compare to manually gated naïve CD8 cells. e) Cells in B25 are divided into two groups: CCR7+ (expression of CCR7 > 1) and CCR7- (expression of CCR7<1). The boxplots show that difference of cell percentage between old and young donors in B25 is driven by CCR7+ cells.

**S7 Fig. Ilustration of box B55 formed by CD4 and CD3.** a) Position of box B55. Cells in B55 express the highest 20 % of both CD3 and CD4. Hence they might be CD4 T cells. b) Percentage of cells in B55 is higher in old donors. c) Scatter plot of mean flourescent intensity (MFI) for all donors and all markers. It suggests cells in B55 might be CD8-, CCR7+ and CD45RA+.

**S8 Fig. Ilustration of box B22 formed by CD4 and CD3.** a) Position of box B22. b) Percentage of cells in B55 is higher in old donors. c) Scatter plot of mean flourescent intensity (MFI) for all donors and all markers. It suggests cells in B22 might be CD11b+, CD14+ and CD45RA+.

**S9 Fig. Ilustration of manual gating strategy for naïve and memory CD8 T cells.**

**S10 Fig.** a) Overlay of cells in B55 on manually gated CD8 naïve and memory cell types for one donor. b) Boxplot of manually gated naïve CD8 cell percentage in live cells.

**S11 Fig. Ilustration of box B51 formed by CD8 and CCR7 (CD8high CCR7low).** a) Position of box B51. b) Boxplot of cell percentage in B51 between young and old donors. c) Scatter plot of mean flourescent intensity (MFI) for all donors and all markers. d & e) MFI of CD45RA vs MFI of CCR7 for cells in B51, naïve and memroy CD8 T cells. Each symbol shows a donor (young donors in d and old donors in e), vertical and horizontal errorbars show standard deviation of CCR7 and CD45RA intensity respectively.

**S1 List. Markers measured in CD4 vs CD8 dataset**

